# Convergent expansions of keystone gene families drive metabolic innovation in a major eukaryotic clade

**DOI:** 10.1101/2024.07.22.604484

**Authors:** Kyle T. David, Joshua G. Schraiber, Johnathan G. Crandall, Abigal L. Labella, Dana A. Opulente, Marie-Claire Harrison, John F. Wolters, Xiaofan Zhou, Xing-Xing Shen, Marizeth Groenewald, Chris Todd Hittinger, Matt Pennell, Antonis Rokas

## Abstract

Many remarkable innovations have repeatedly occurred across vast evolutionary distances. When convergent traits emerge on the tree of life, they are sometimes driven by the same underlying gene families, while other times many different gene families are involved. Conversely, a gene family may be repeatedly recruited for a single trait or many different traits. To understand the general rules governing convergence at both genomic and phenotypic levels, we systematically tested associations between 56 binary metabolic traits and gene count in 14,710 gene families from 993 species of *Saccharomycotina* yeasts. Using a recently developed phylogenetic approach that reduces spurious correlations, we discovered that gene family expansion and contraction was significantly linked to trait gain and loss in 45/56 (80%) of traits. While 601/746 (81%) of significant gene families were associated with only one trait, we also identified several ‘keystone’ gene families that were significantly associated with up to 13/56 (23%) of all traits. These results indicate that metabolic innovations in yeasts are governed by a narrow set of major genetic elements and mechanisms.

## Introduction

The repeated occurrences of biological innovations have long been used to provide strong evidence for the power of natural selection and predictability of evolution^1^. However, the extent to which convergent traits are achieved *via* the same genetic elements (parallelism) remains a longstanding question^2–5^ (Fig. 1). For example, animal eyes are a classic example of convergent evolution, having emerged at least 40 times across Metazoa^6^. Due to the vast evolutionary distances and morphological variation found across eyes, they were long assumed to have evolved *via* an equally diverse number of genetic pathways^7^. However, it is now established that development of all animal eyes is globally controlled by a *PAX*6 homolog^8,9^, independently recruited many times to achieve the same trait in a remarkable case of parallelism. In contrast, antifreeze proteins, another classic case of convergent evolution, have repeatedly evolved in a non-parallel fashion from diverse genomic origins. Even proteins with virtually identical structures and sequences have evolved from at least three distinct gene families^10^.

**Figure 1.**
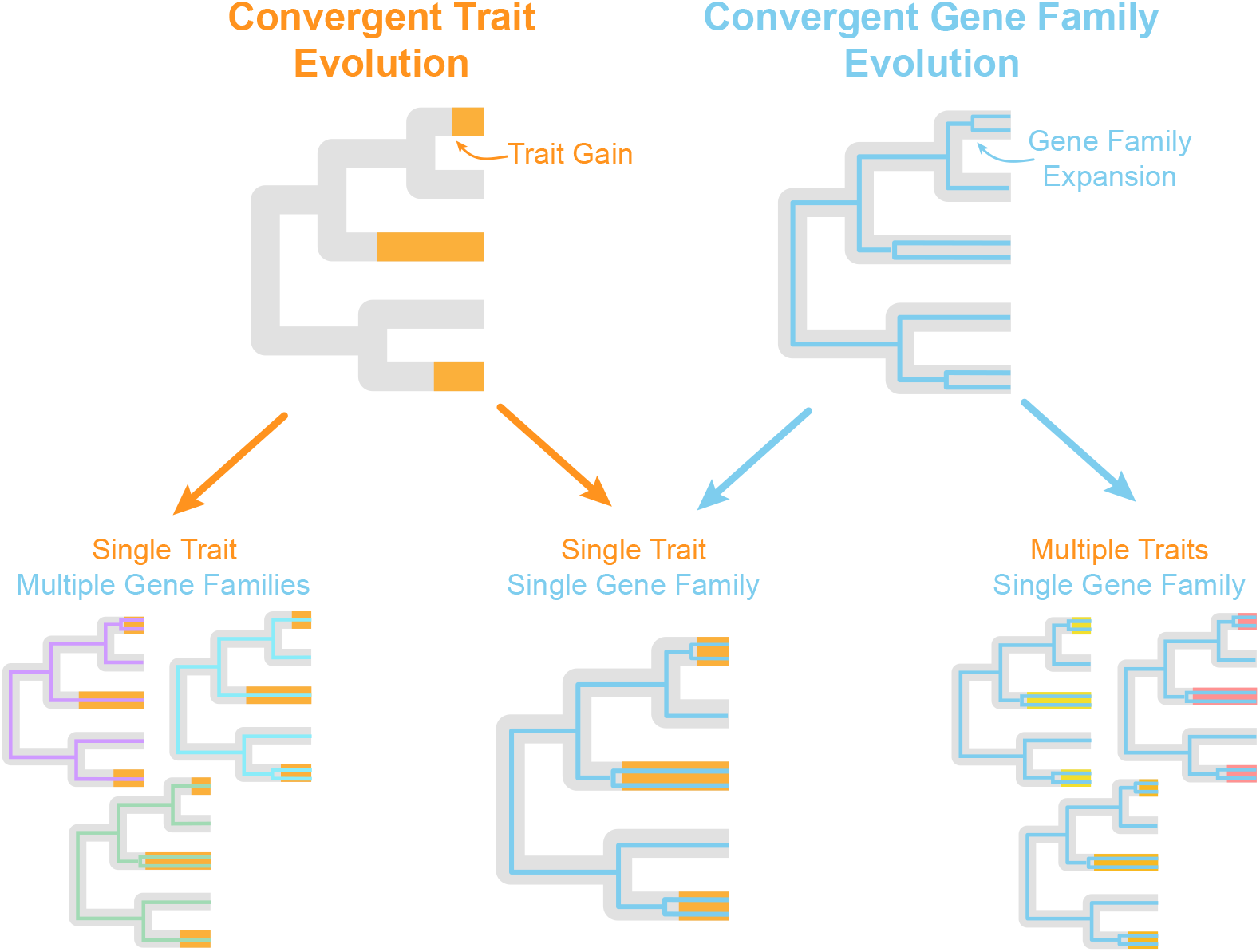
Models of convergent evolution. When convergent traits occur (top left), they may be the result of multiple (nonparallel; bottom left) or single (parallel; bottom middle) underlying genetic elements. Likewise, convergent genetic changes (top right) may have effects on single (bottom middle) or multiple (bottom right) traits.

Image-forming eyes and antifreeze proteins are highly specialized traits associated with specific functions. More general traits that are essential to the basic maintenance of an organism are expected to be more conserved and experience stronger purifying selection, such that convergence may be limited^5,7,11^. However even for fundamental housekeeping processes, convergent solutions occur. For example, oxygen transport has evolved many times through both parallel and nonparallel means^7^: the protein hemerythrin has been recruited at least four times across three animal phyla^12^ for oxygen transport, and is itself analogous to other convergently evolving O_2_-binding protein families like the hemocyanins and hemoglobins^13^.

Examples of nonparallel and (especially) parallel convergent evolution across deeper evolutionary distances, such as those described above, are few and far between. Much of the previous work studying convergence at genomic and phenotypic levels has focused on relatively recent changes, identifying convergent patterns in single genes or even nucleotides across populations^14^. However, it is difficult to distinguish true convergence from shared ancestral genetic variation on microevolutionary scales^11,15,16^.

Gene family evolutionary events such as gene duplication, gene loss, and horizontal gene transfer are often invoked in trait evolution and innovation^17,18^. However, support for causal relationships is often weak, as evolutionary innovations often evolve only once^19^, leaving no degrees of freedom to statistically test gene family – trait evolution associations^20^. To overcome these issues, we selected 56 metabolic traits, each of which evolved dozens to thousands of times across 400 million years of evolution, providing unprecedented statistical power for investigating convergent evolutionary events. These 56 metabolic traits have been experimentally tested^21,22^ for 993 species of *Saccharomycotina* yeasts (“yeasts” hereafter). Yeasts are able to metabolize alcohols, ketones, organic acids and more, which has enabled them to colonize virtually every continent and biome on the planet^23,24^. The spectacular diversity of yeast metabolism has not gone unnoticed by humans. In addition to the genus *Saccharomyces*, whose metabolisms underwrite the baking, brewing, and winemaking industries, many yeasts such as *Yarrowia lipolytica, Lipomyces starkeyi*, and *Komagataella (Pichia) pastoris* have unique metabolisms that are exploited for technological and industrial applications^25–27^. By sampling genomes from ∼80% of described *Saccharomycotina* species and traits from across the metabolic spectrum we were able to systematically quantify the number of evolutionary paths for trait innovation across a wide swath of genetic and phenotypic diversity.

### Parallel gene family expansion undergirds metabolic diversity in yeasts

Metabolic variation in *Saccharomycotina* is extensive, from specialist species able to metabolize one or two compounds to extreme generalists found growing on 47/56 (84%) of tested substrates. Genetic variation is also broad; a family of RNA-directed DNA polymerases (K00986) is absent or represented by a single gene in certain species but can have as many as 290 genes in others. To test associations between gene family size and metabolic ability, we ran a modified phylogenetic logistic regression model for each combination of traits and gene families (N = 823,760 tests). We identified 746/14,710 (5%) significant (FDR<10^−6^) families. These families were significantly (FDR<0.05) enriched in four KEGG pathways, all of which belong to the metabolism category, including sucrose metabolism and metabolism in diverse environments (Table S1). 45/56 (80%) traits were significantly associated with at least one (on average 15) gene families, wherein the size of the family repeatedly predicted metabolic ability across lineages (Fig. 2). This relationship was positive in 697/746 (93%) families, strongly implicating gene family expansion as an engine of parallel metabolic innovation in yeasts. To investigate such an engine, we estimated the evolutionary history of the most highly significant association between a trait (raffinose) and gene family (*SUC*) in our analyses. In *S. cerevisiae, SUC2* encodes the enzyme that cleaves the glycosidic link in raffinose, producing fructose and melibiose. As expected, we found that gene duplication and horizontal transfer events within the family map closely with trait gains across the yeast phylogeny (Fig. 2).

**Figure 2.**
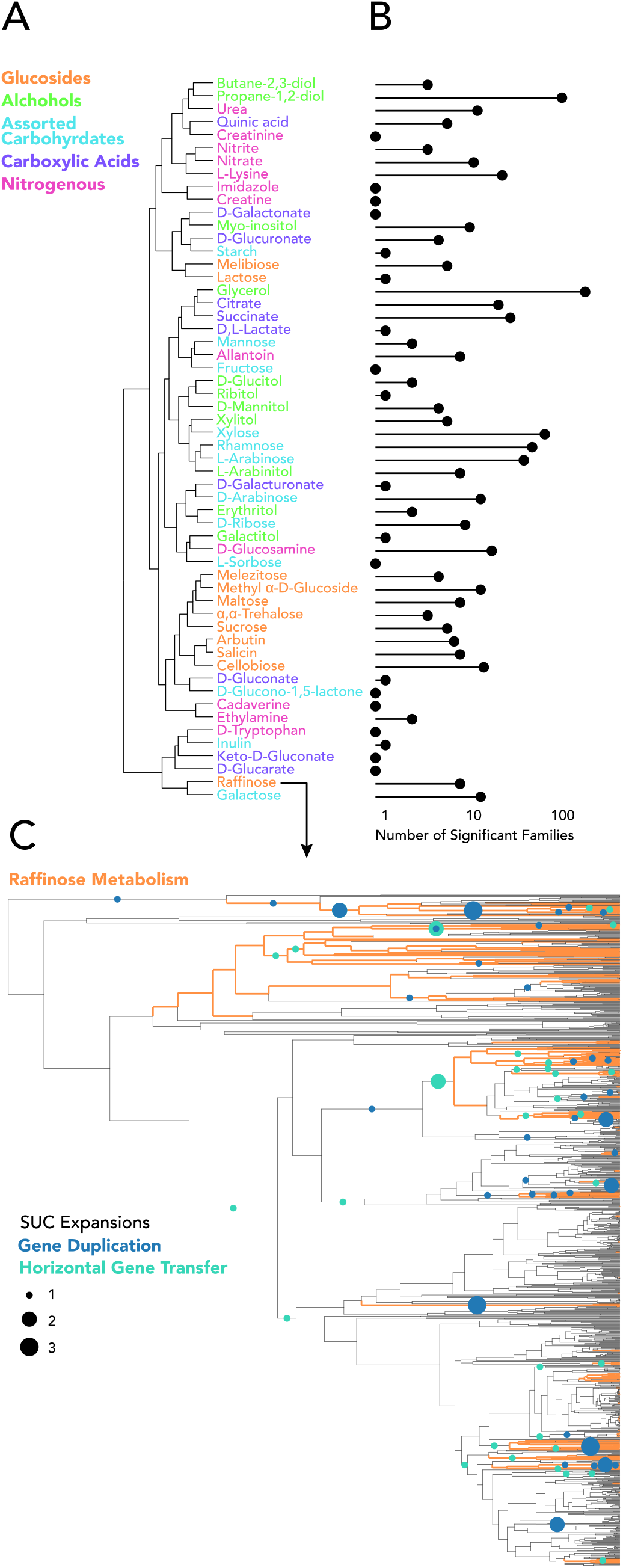
Gene family expansion and contraction was significantly linked to trait gain and loss in 80% of metabolic traits. A) All 56 metabolic traits examined by this study, clustered by gene family affinity. Traits are clustered by z-scores across families, closer traits share more similar correlations with the same gene families. B) Number of gene families that are significantly associated with each metabolic trait. C) Example of a convergent trait (raffinose metabolism), driven by convergent expansions within the same gene family (*SUC*). The ability to metabolize raffinose is mapped alongside gene duplication and horizontal gene transfer events of the *SUC* gene family on the yeast species phylogeny.

The extent of gene family parallelism also reflects the biochemical properties of the metabolic substrates examined. For example, 8/11 (72%) traits without evidence for parallelism were for carboxylic acids or nitrogenous substrates. By contrast, all alcohols and glucosides had strong evidence of parallelism. This result suggests that the number of evolutionary solutions to metabolize the latter may be more limited than the former. To further compare patterns of gene family usage across traits, we performed hierarchical clustering of z-scores for each regression, meaning traits that clustered closer together shared more associations with the same gene families (Fig. 2). We found that closely related compounds (e.g. nitrate and nitrite, isomers butane-2,3-diol and propane-1,2-diol, xylose and its alcohol derivative xylitol) were often reciprocal nearest neighbors. Shared usage of gene families between similar chemical substrates implies that certain families may be repeatedly recruited for the convergent evolution of similar but distinct traits.

### Keystone gene families drive convergent evolution of multiple traits

Our results raise the hypothesis that the same gene families may be repeatedly co-opted in the evolution of multiple convergent traits. To formally test this hypothesis and measure the extent of shared usage, we examined how many traits were significantly associated with each gene family (Fig. 3). As expected, the majority (601/746) of significant gene families were only associated with one specific trait. However, we also identified 145/746 (19%) gene families associated with multiple traits. Of particular note was a gene family encoding α-glucoside transporters (*AGT*), which was significantly associated with the convergent evolution of 13 different traits, almost twice as many as any other family. In *S. cerevisiae*, members of this family are known to encode a generalist transporter (Agt1) and three more specialized transporters (Mal31, Mph2, and Mph3) ^28,29^. We find strong evidence this family has been repeatedly recruited to transport all known Agt1 substrates across the evolution of yeasts, as well as most glucosides (save for lactose and raffinose), in addition the nitrogen sources urea and the monosaccharides arabinose, rhamnose, and xylose. Of the other 14 ‘keystone’ families (defined as those that exhibit significant associations with the convergent evolution of at least 4 different traits), all encoded either other transporters or enzymes that catalyze decomposition reactions such as hydrolases and lyases (Table S2). The repeated recruitment of the same gene family for similar but distinct traits suggests that these families provide important functions across multiple metabolic pathways. Just as keystone species demonstrate an outsized impact on their ecological communities^30^, these keystone families appear to have been instrumental in the evolution of metabolism across a wide variety of chemical substrates.

**Figure 3.**
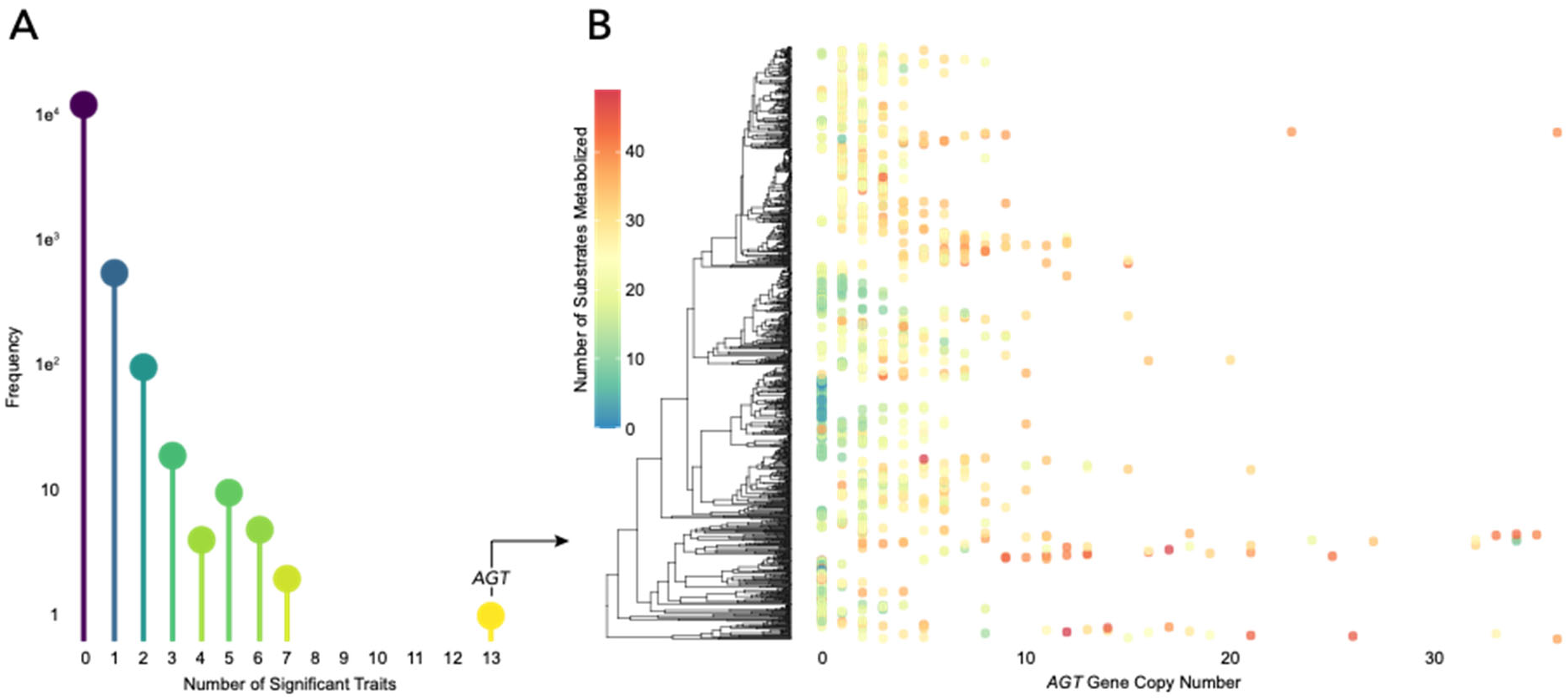
Keystone gene families influence innovation of multiple metabolic traits. A) Histogram of the number of traits significantly associated with each of 14,710 gene families. Of the 746 families with a significant relationship, most (81%) are associated with just one trait, with the remaining 19% associated with two or more traits. Furthermore, a minority of keystone families show significant effects on several (≥ 4) traits. B) An extreme example of a keystone gene family encoding α-glucoside transporters (*AGT*). Size of the *AGT* gene family mapped onto a species phylogeny, colored by the total number of substrates metabolized by each species. On average, the *AGT* family is larger in generalist species than in specialist species.

## Conclusion

Using genomes and metabolic trait data for 56 substrates from 993 *Saccharomycotina* yeasts, we found that the convergent evolution of metabolic traits coincides with convergent gene family evolution in 45/56 (80%) of cases, showing widespread evidence of parallelism on the gene family level. Perhaps more remarkably, we also found that convergent evolution in certain gene families coincides with the convergent evolution of *multiple* traits. The incidence of parallelism in convergent evolution is expected to decrease with divergence time, with limited examples beyond the species level^14,31^. However, we find extensive parallel evolution across 400 million years of *Saccharomycotina* evolution, who possess roughly as much genetic diversity as the plant and animal kingdoms^32^. We further identify hundreds of gene families that are significantly linked to metabolic innovations across *Saccharomycotina*, offering putative targets for genetic engineering. Predicting evolution is of particular interest for this clade, whose metabolisms are harnessed by humans for a slew of purposes across medical, scientific, and industrial fields^23^.

Gene family expansions are often expected to lead to phenotypic innovation, either through neofunctionalization of duplicate genes^17^ or sharing functions with other species through horizontal transfer^33^. For example, whole genome duplication events have been found at several critical junctures in macroevolution, preceding adaptive radiations and mass extinction events^34^. In yeasts, the aerobic fermentative abilities of the *Saccharomyces* genus have long been attributed to a whole genome duplication^35^. However, as evolutionary innovations are typically rare, sufficient sample sizes to identify these drivers are often absent. The high numbers of convergent events in both trait and gene family evolution provided by this study offer exceptional statistical power to address these questions. We find strong evidence for the importance of gene family expansion in macroevolution, with variation in gene family size tightly linked to metabolic ability in 45/56 (80%) of traits tested. The repeated association between gene family expansion events and trait gains across traits and lineages suggests that gene duplication and horizontal transfer are a common pathway to evolutionary innovation.

Our discovery that convergently evolved metabolic traits are often driven by the same underlying gene families has important implications for the predictability of evolution. We found strong evidence for phenotypic and genomic convergence in 80% of traits tested, suggesting parallel evolution across deep timescales (such as in *PAX*6 and the development of animal eyes) may be far more common than previously believed. Remarkably, we have also discovered certain gene families involved in the convergence of multiple traits, with the repeated evolution of over a dozen diverse metabolic traits seemingly contingent on gene gains and losses within a single family of transporters. Taken together, these results invoke a more deterministic view of innovation^2,5,36^, wherein novel traits are gained through a narrow set of possible genetic elements and mechanisms.

## Methods

### Genomic dataset

Genomes and the species phylogeny were obtained from Opulente *et al*. 2024^21^. Briefly, 1,154 yeast species were paired-end sequenced on an Illumina HiSeq 25000, and genes were predicted with the BRAKER^37^ v2.1.6 pipeline. After aligning, a species phylogeny was inferred with IQ-TREE^38^ v2.0.7 using the general amino-acid substitution matrix^39^ with four gamma discretized rate categories. The phylogeny was then time-calibrated using the RelTime method implemented in MEGA7^40^ using calibration points from Shen *et al*. 2018^32^. Additional information, as well as all genomes, alignments, and phylogenies can be found at the original publication^21^. OrthoFinder^41^ v3.0 was run under default parameters on the 1,154 genomes, resulting in 72,381 gene families^18^ which were then filtered to 14,710 to include only gene families with at least 10 species represented. Following Opulente *et al*. 2024^21^, gene models were annotated with KEGG orthologs^42^ using KofamScan^43^. Enrichment analysis was performed using the enrichKEGG() function in the R package clusterProfiler^44^ v.4.10.

### Trait dataset

Metabolic traits for each species were sourced from Harrison *et al*. 2024^22^, and supplemented with experimental assays from Opulente *et al*. 2024^21^. Metabolic traits were considered present if cultures were able to grow on 96-well plates containing the carbon or nitrogen source in minimal media. Traits and genomes were merged based on the most recently available taxonomy^45^ hosted on the MycoBank database^46^. Traits were retained if they had no more than 50% missing data across the 993 species. Two additional metabolic traits were further removed: growth on glucose, which was present in every species, and growth in the absence of carbon, which is not directly comparable with positive substrate-specific metabolic traits. The final trait matrix had 19% missing data across 46 carbon and 10 nitrogen substrates (Fig. S1). Evolution of each trait was modeled using a stochastic character mapping approach implemented in RevBayes^47^ v1.2.2 using two hidden states. Models were run for 1000 generations across two chains.

### Phylogenetic analyses

To assess the effect of gene family size on trait evolution we used a phylogenetic logistic regression as implemented in the phylolm^48^ v2.6.2 R^49^ v4.3.2 package using the maximum penalized likelihood estimation method. A square root transformation was applied to gene family size to reflect the expected diminishing contribution of additional genes in large families.

When studying patterns of convergence care must be taken to distinguish truly independent events from synapomorphies shared by common descent^7^. Conventional phylogenetic comparative methods have been shown to have trouble distinguishing between these scenarios, attributing significance for spurious correlations even for single unreplicated events^20,50^. It was recently shown that many approaches in phylogenetic comparative methods and statistical genetics represent special cases of the same general model^51^. Leveraging this discovery, we adapt a common strategy in genome-wide association studies shown to reduce these types of spurious correlations^20,51^ by including eigenvectors of the phylogenetic variance-covariance matrix as fixed effects in our phylogenetic logistic regression model. In our analysis we selected the first two leading eigenvectors, which together contain over 50% of shared variance across the phylogeny. As expected, these eigenvectors explain the greatest variance in traits with the fewest number of transitions, supporting the idea that they reduce false positives resulting from shared ancestral events (Fig. S2). To further reduce the possibility of false positives and correct for multiple tests, we calculated the false discovery rate^52^ adopting a conservative alpha value of 10^−6^. Evolution of the *fructan beta-fructosidase* family was estimated with GeneRax^53^ v2.0.4, a species-gene tree reconciliation program, using a maximum subtree prune and regraft distance of 3. The starting gene tree was inferred using IQ-TREE^38^ v2.2.2 and both initial and reconciled gene trees used 10 FreeRate^54,55^ categories while also allowing for invariant sites, the model parameters with the highest Bayesian information criterion according to ModelFinder^56^.

## Data Availability

All underlying data and code will be made available upon publication.

## Acknowledgements

This work was performed using resources contained within the Advanced Computing Center for research and Education at Vanderbilt University in Nashville, TN.

## References

1. Smith, S. D., Pennell, M. W., Dunn, C. W. & Edwards, S. V. Phylogenetics is the New Genetics (for Most of Biodiversity). Trends in Ecology & Evolution 35, 415–425 (2020).

2. Oakley, T. Building, Maintaining, and (Re-) Deploying Genetic Toolkits during Convergent Evolution. (2024). at <https://www.preprints.org/manuscript/202404.1289>

3. Sackton, T. B., Grayson, P., Cloutier, A., Hu, Z., Liu, J. S., Wheeler, N. E., Gardner, P. P., Clarke, J. A., Baker, A. J., Clamp, M. & Edwards, S. V. Convergent regulatory evolution and loss of flight in paleognathous birds. Science 364, 74–78 (2019).

4. Losos, J. B. Convergence, Adaptation, and Constraint. Evolution 65, 1827–1840 (2011).

5. Wake, D. B., Wake, M. H. & Specht, C. D. Homoplasy: From Detecting Pattern to Determining Process and Mechanism of Evolution. Science 331, 1032–1035 (2011).

6. Land, M. F. & Fernald, R. D. The Evolution of Eyes. Annu. Rev. Neurosci. 15, 1–29 (1992).

7. McGhee, G. R. Convergent evolution: limited forms most beautiful. (MIT press, 2011).

8. Tomarev, S. I., Callaerts, P., Kos, L., Zinovieva, R., Halder, G., Gehring, W. & Piatigorsky, J. Squid Pax-6 and eye development. Proc. Natl. Acad. Sci. U.S.A. 94, 2421–2426 (1997).

9. Halder, G., Callaerts, P. & Gehring, W. J. Induction of Ectopic Eyes by Targeted Expression of the eyeless Gene in Drosophila. Science 267, 1788–1792 (1995).

10. Rives, N., Lamba, V., Cheng, C.-H.C. & Zhuang, X. Diverse origins of near-identical antifreeze proteins in unrelated fish lineages provide insights into evolutionary mechanisms of new gene birth and protein sequence convergence. Preprint at 10.1101/2024.03.12.584730 (2024)

11. Stern, D. L. The genetic causes of convergent evolution. Nat Rev Genet 14, 751–764 (2013).

12. Bailly, X., Vanin, S., Chabasse, C., Mizuguchi, K. & Vinogradov, S. N. A phylogenomic profile of hemerythrins, the nonheme diiron binding respiratory proteins. BMC Evol Biol 8, 244 (2008).

13. Coates, C. J. & Decker, H. Immunological properties of oxygen-transport proteins: hemoglobin, hemocyanin and hemerythrin. Cell. Mol. Life Sci. 74, 293–317 (2017).

14. Bohutínská, M. & Peichel, C. L. Divergence time shapes gene reuse during repeated adaptation. Trends in Ecology & Evolution 39, 396–407 (2024).

15. Soria-Carrasco, V., Gompert, Z., Comeault, A. A., Farkas, T. E., Parchman, T. L., Johnston, J. S., Buerkle, C. A., Feder, J. L., Bast, J., Schwander, T., Egan, S. P., Crespi, B. J. & Nosil, P. Stick Insect Genomes Reveal Natural Selection’s Role in Parallel Speciation. Science 344, 738–742 (2014).

16. Colosimo, P. F., Hosemann, K. E., Balabhadra, S., Villarreal, G., Dickson, M., Grimwood, J., Schmutz, J., Myers, R. M., Schluter, D. & Kingsley, D. M. Widespread Parallel Evolution in Sticklebacks by Repeated Fixation of Ectodysplasin Alleles. Science 307, 1928– 1933 (2005).

17. Ohno, S. Evolution by gene duplication. (Springer Science & Business Media, 1970).

18. Feng, B., Li, Y., Liu, H., Steenwyk, J. L., David, K. T., Tian, X., Xu, B., Gonçalves, C., Opulente, D. A., LaBella, A. L., Harrison, M.-C., Wolters, J. F., Shao, S., Chen, Z., Fisher, K. J., Groenewald, M., Hittinger, C. T., Shen, X.-X., Rokas, A., Zhou, X. & Li, Y. Unique trajectory of gene family evolution from genomic analysis of nearly all known species in an ancient yeast lineage. 2024.06.05.597512 Preprint at 10.1101/2024.06.05.597512 (2024)

19. Vermeij, G. J. Historical contingency and the purported uniqueness of evolutionary innovations. Proc. Natl. Acad. Sci. U.S.A. 103, 1804–1809 (2006).

20. Uyeda, J. C., Zenil-Ferguson, R. & Pennell, M. W. Rethinking phylogenetic comparative methods. Systematic Biology 67, 1091–1109 (2018).

21. Opulente, D. A., LaBella, A. L., Harrison, M.-C., Wolters, J. F., Liu, C., Li, Y., Kominek, J., Steenwyk, J. L., Stoneman, H. R., VanDenAvond, J., Miller, C. R., Langdon, Q. K., Silva, M., Gonçalves, C., Ubbelohde, E. J., Li, Y., Buh, K. V., Jarzyna, M., Haase, M. A. B., Rosa, C. A., ČCadež, N., Libkind, D., DeVirgilio, J. H., Hulfachor, A. B., Kurtzman, C. P., Sampaio, J. P., Gonçalves, P., Zhou, X., Shen, X.-X., Groenewald, M., Rokas, A. & Hittinger, C. T. Genomic factors shape carbon and nitrogen metabolic niche breadth across Saccharomycotina yeasts. Science 384, eadj4503 (2024).

22. Harrison, M.-C., Ubbelohde, E. J., LaBella, A. L., Opulente, D. A., Wolters, J. F., Zhou, X., Shen, X.-X., Groenewald, M., Hittinger, C. T. & Rokas, A. Machine learning enables identification of an alternative yeast galactose utilization pathway. Proc. Natl. Acad. Sci. U.S.A. 121, e2315314121 (2024).

23. Kurtzman, C. P., Fell, J. W. & Boekhout, T. The yeasts: a taxonomic study. (Elsevier, 2011).

24. David, K. T., Harrison, M.-C., Opulente, D. A., LaBella, A. L., Wolters, J. F., Zhou, X., Shen, X.-X., Groenewald, M., Pennell, M., Hittinger, C. T. & Rokas, A. Saccharomycotina yeasts defy long-standing macroecological patterns. Proc. Natl. Acad. Sci. U.S.A. 121, e2316031121 (2024).

25. Spagnuolo, M., Yaguchi, A. & Blenner, M. Oleaginous yeast for biofuel and oleochemical production. Current Opinion in Biotechnology 57, 73–81 (2019).

26. Gassler, T., Sauer, M., Gasser, B., Egermeier, M., Troyer, C., Causon, T., Hann, S., Mattanovich, D. & Steiger, M. G. The industrial yeast Pichia pastoris is converted from a heterotroph into an autotroph capable of growth on CO2. Nat Biotechnol 38, 210–216 (2020).

27. Srinivasan, P. & Smolke, C. D. Biosynthesis of medicinal tropane alkaloids in yeast. Nature 585, 614–619 (2020).

28. Han, E., Cotty, F., Sottas, C., Jiang, H. & Michels, C. A. Characterization of AGT1 encoding a general α-glucoside transporter from Saccharomyces. Molecular Microbiology 17, 1093–1107 (1995).

29. Stambuk, B. U., da Silva, M. A., Panek, A. D. & de Araujo, P. S. Active α-glucoside transport in Saccharomyces cerevisiae. FEMS microbiology letters 170, 105–110 (1999).

30. Paine, R. T. A Note on Trophic Complexity and Community Stability. The American Naturalist 103, 91–93 (1969).

31. Yeaman, S., Hodgins, K. A., Lotterhos, K. E., Suren, H., Nadeau, S., Degner, J. C., Nurkowski, K. A., Smets, P., Wang, T., Gray, L. K., Liepe, K. J., Hamann, A., Holliday, J. A., Whitlock, M. C., Rieseberg, L. H. & Aitken, S. N. Convergent local adaptation to climate in distantly related conifers. Science 353, 1431–1433 (2016).

32. Shen, X.-X., Opulente, D. A., Kominek, J., Zhou, X., Steenwyk, J. L., Buh, K. V., Haase, M. A. B., Wisecaver, J. H., Wang, M., Doering, D. T., Boudouris, J. T., Schneider, R. M., Langdon, Q. K., Ohkuma, M., Endoh, R., Takashima, M., Manabe, R., Čadež, N., Libkind, D., Rosa, C. A., DeVirgilio, J., Hulfachor, A. B., Groenewald, M., Kurtzman, C. P., Hittinger, C. T. & Rokas, A. Tempo and Mode of Genome Evolution in the Budding Yeast Subphylum. Cell 175, 1533-1545.e20 (2018).

33. Etten, J. V. & Bhattacharya, D. Horizontal Gene Transfer in Eukaryotes: Not if, but How Much? Trends in Genetics 36, 915–925 (2020).

34. Van de Peer, Y., Mizrachi, E. & Marchal, K. The evolutionary significance of polyploidy. Nature Reviews Genetics 18, 411 (2017).

35. Wolfe, K. H. & Shields, D. C. Molecular evidence for an ancient duplication of the entire yeast genome. Nature 387, 708–713 (1997).

36. Losos, J. B., Jackman, T. R., Larson, A., Queiroz, K.de & Rodríguez-Schettino, L. Contingency and Determinism in Replicated Adaptive Radiations of Island Lizards. Science 279, 2115–2118 (1998).

37. Brůna, T., Hoff, K. J., Lomsadze, A., Stanke, M. & Borodovsky, M. BRAKER2: automatic eukaryotic genome annotation with GeneMark-EP+ and AUGUSTUS supported by a protein database. NAR Genomics and Bioinformatics 3, lqaa108 (2021).

38. Nguyen, L.-T., Schmidt, H. A., Von Haeseler, A. & Minh, B. Q. IQ-TREE: a fast and effective stochastic algorithm for estimating maximum-likelihood phylogenies. Molecular biology and evolution 32, 268–274 (2015).

39. Le, S. Q. & Gascuel, O. An improved general amino acid replacement matrix. Molecular biology and evolution 25, 1307–1320 (2008).

40. Kumar, S., Stecher, G. & Tamura, K. MEGA7: Molecular Evolutionary Genetics Analysis Version 7.0 for Bigger Datasets. Molecular Biology and Evolution 33, 1870–1874 (2016).

41. Emms, D. M. & Kelly, S. OrthoFinder: phylogenetic orthology inference for comparative genomics. Genome biology 20, 1–14 (2019).

42. Kanehisa, M. & Goto, S. KEGG: kyoto encyclopedia of genes and genomes. Nucleic acids research 28, 27–30 (2000).

43. Aramaki, T., Blanc-Mathieu, R., Endo, H., Ohkubo, K., Kanehisa, M., Goto, S. & Ogata, H. KofamKOALA: KEGG Ortholog assignment based on profile HMM and adaptive score threshold. Bioinformatics 36, 2251–2252 (2020).

44. Wu, T., Hu, E., Xu, S., Chen, M., Guo, P., Dai, Z., Feng, T., Zhou, L., Tang, W. & Zhan, L. I. clusterProfiler 4.0: A universal enrichment tool for interpreting omics data. The innovation 2, (2021).

45. Groenewald, M., Hittinger, C. T., Bensch, K., Opulente, D. A., Shen, X.-X., Li, Y., Liu, C., LaBella, A. L., Zhou, X. & Limtong, S. A genome-informed higher rank classification of the biotechnologically important fungal subphylum Saccharomycotina. Studies in Mycology (2023).

46. Crous, P. W., Gams, W., Stalpers, J. A., Robert, V. & Stegehuis, G. MycoBank: an online initiative to launch mycology into the 21st century. Studies in mycology 50, 19–22 (2004).

47. Höhna, S., Landis, M. J., Heath, T. A., Boussau, B., Lartillot, N., Moore, B. R., Huelsenbeck, J. P. & Ronquist, F. RevBayes: Bayesian phylogenetic inference using graphical models and an interactive model-specification language. Systematic Biology 65, 726–736 (2016).

48. Ho, L. S. T., Ane, C., Lachlan, R., Tarpinian, K., Feldman, R., Yu, Q., van der Bijl, W., Maspons, J., Vos, R. & Ho, M. L. S. T. Package ‘phylolm’. See http://cran.r-project.org/web/packages/phylolm/index.html (accessed February 2018) (2016).

49. Team, R. C. R: A Language and Environment for Statistical Computing. (2017). At <https://www.R-project.org/>

50. Maddison, W. P. & FitzJohn, R. G. The Unsolved Challenge to Phylogenetic Correlation Tests for Categorical Characters. Systematic Biology 64, 127–136 (2015).

51. Schraiber, J. G., Edge, M. D. & Pennell, M. Unifying approaches from statistical genetics and phylogenetics for mapping phenotypes in structured populations. bioRxiv 2024–02 (2024).

52. Benjamini, Y. & Hochberg, Y. Controlling the false discovery rate: a practical and powerful approach to multiple testing. Journal of the Royal statistical society: series B (Methodological) 57, 289–300 (1995).

53. Morel, B., Kozlov, A. M., Stamatakis, A. & Szöll\Hosi, G. J. GeneRax: a tool for species-tree-aware maximum likelihood-based gene family tree inference under gene duplication, transfer, and loss. Molecular biology and evolution 37, 2763–2774 (2020).

54. Yang, Z. A space-time process model for the evolution of DNA sequences. Genetics 139, 993–1005 (1995).

55. Soubrier, J., Steel, M., Lee, M. S. Y., Der Sarkissian, C., Guindon, S., Ho, S. Y. W. & Cooper, A. The Influence of Rate Heterogeneity among Sites on the Time Dependence of Molecular Rates. Molecular Biology and Evolution 29, 3345–3358 (2012).

56. Kalyaanamoorthy, S., Minh, B. Q., Wong, T. K., von Haeseler, A. & Jermiin, L. S. ModelFinder: fast model selection for accurate phylogenetic estimates. Nature methods 14, 587 (2017).

