## Supplementary Figures and Tables for "Convergent expansions of keystone gene families drive metabolic innovation in a major eukaryotic clade"

**Supplemental Material**

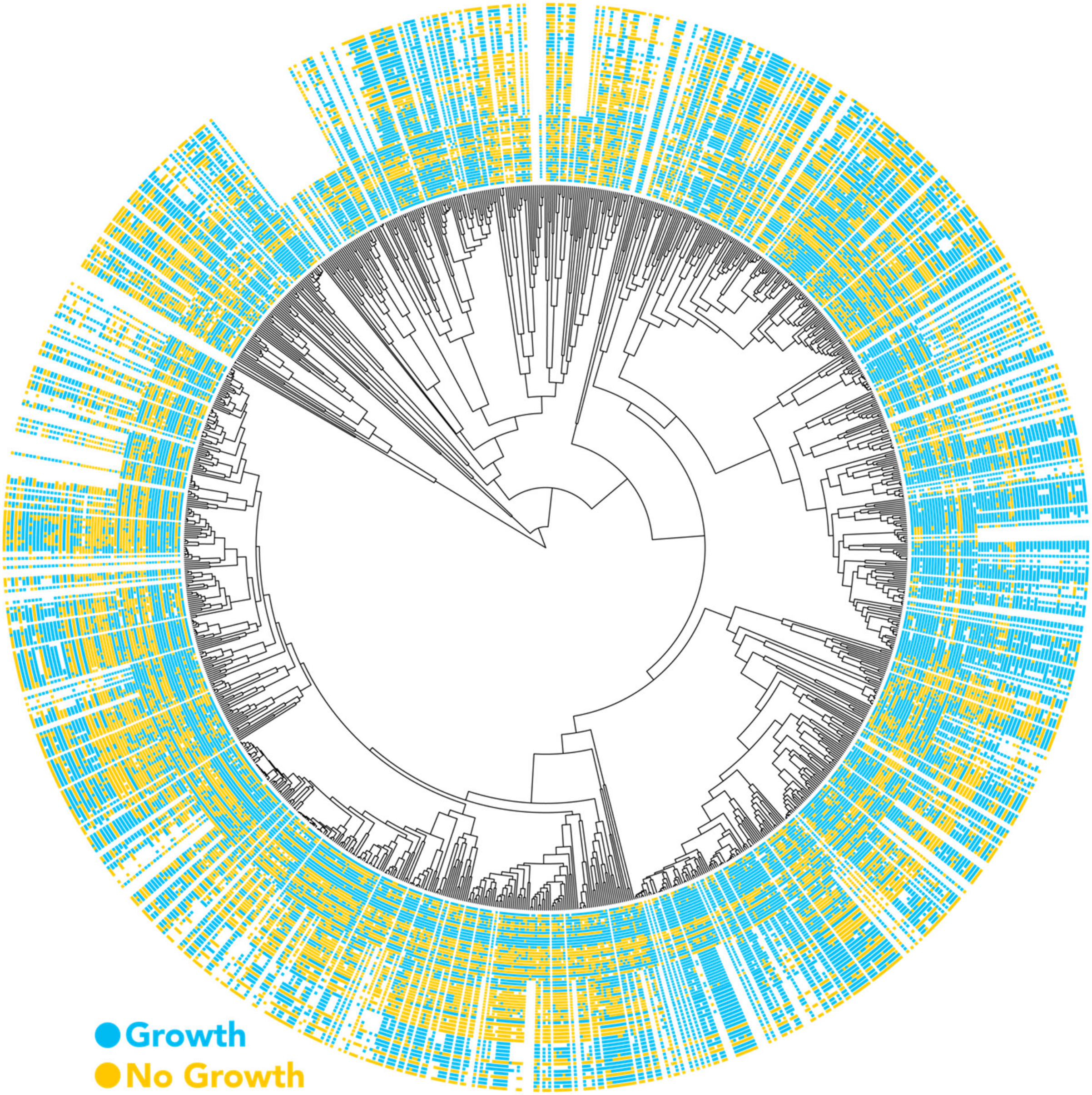


**Figure S1.** Visualization of data occupancy and phylogenetic sampling. Rows indicate whether or not a trait is present, absent, or missing for a given species, mapped onto the yeast species phylogeny. The full matrix and underlying data will be made available upon publication.

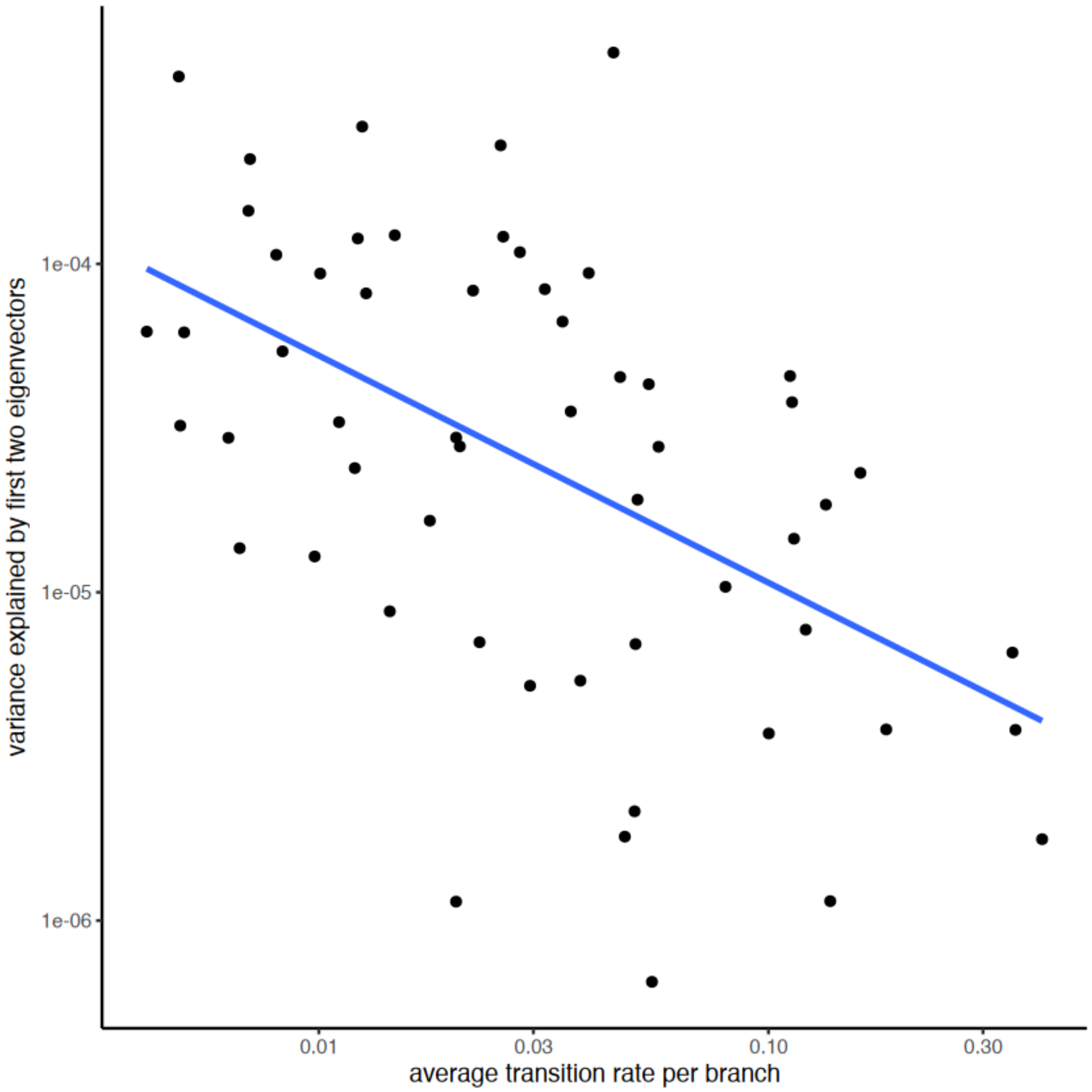


**Figure S2.** Transition rate vs. variance explained by phylogenetic covariance for each of the 56 metabolic traits. Phylogenetic variance is more informative for slowly evolving traits. As expected, traits that evolve more rapidly are less well explained by phylogenetic covariance, indicating the absence of phylogenetic inertia.

**Table S1.** KEGG enrichment analysis results of all significant gene families.

| Category | ID | Description | GeneRatio | p.adjust |
|---|---|---|---|---|
| **Metabolism** | ko00052 | Galactose metabolism | 11/364 | 0.0003647 |
| **Metabolism** | ko00051 | Fructose and mannose metabolism | 14/364 | 0.00084482 |
| **Metabolism** | ko01120 | Microbial metabolism in diverse environments | 52/364 | 0.01517963 |
| **Metabolism** | ko00040 | Pentose and glucuronate interconversions | 9/364 | 0.03509469 |

**Table S2.** KEGG annotations of keystone gene families associated with more than three traits. When multiple annotations were found within the same family, the most common one was used.

| OG | KO | name | class | significant traits |
|---|---|---|---|---|
| **OG0000047** | K08141 | MFS transporter, SP family, general alpha glucoside:H+ symporter | Transporter: sugar transporter | 13 |
| **OG0000277** | K05351 | D-xylulose reductase | Enzyme: Oxidoreductase | 7 |
| **OG0004218** | K01179 | endoglucanase | Enzyme: Hydrolase | 7 |
| **OG0000038** | NA | NA | NA | 6 |
| **OG0000048** | K08139 | MFS transporter, SP family, sugar:H+ symporter | Transporter: sugar transporter | 6 |
| **OG0000141** | K08174 | MFS transporter, FHS family, glucose/mannose:H+ symporter | Transporter: sugar transporter | 6 |
| **OG0000447** | K01182 | oligo-1,6-glucosidase | Enzyme: Hydrolase | 6 |
| **OG0004931** | K18334 | L-fuconate dehydratase | Enzyme: Lyase | 6 |
| **OG0000007** | K08192 | MFS transporter, ACS family, DAL5 transporter family protein | Transporter: organic acid transporter | 5 |
| **OG0000010** | K17742 | sorbose reductase | Enzyme: Oxidoreductase | 5 |
| **OG0000036** | K03327 | MATE family, multidrug and toxin extrusion protein | Transporter: solute carrier | 5 |
| **OG0000261** | K05349 | beta-glucosidase | Enzyme: Hydrolase | 5 |
| **OG0000557** | NA | NA | NA | 5 |
| **OG0000754** | K18339 | 2-keto-3-deoxy-L-rhamnonate aldolase | Enzyme: Lyase | 5 |
| **OG0004213** | NA | NA | NA | 5 |
| **OG0004256** | K01192 | beta-mannosidase | Enzyme: Hydrolase | 5 |
| **OG0004426** | K12661 | L-rhamnonate dehydratase | Enzyme: Lyase | 5 |
| **OG0004764** | K18338 | L-rhamnono-1,4-lactonase | Enzyme: Hydrolase | 5 |
| **OG0000493** | K10256 | omega-6 fatty acid desaturase / acyl-lipid omega-6 desaturase (Delta-12 desaturase) | Enzyme: Oxidoreductase | 4 |
| **OG0000554** | K00306 | sarcosine oxidase / L-pipecolate oxidase | Enzyme: Oxidoreductase | 4 |
| **OG0000796** | K01738 | cysteine synthase | Enzyme: Transferase | 4 |
| **OG0004491** | K01811 | alpha-D-xyloside xylohydrolase | Enzyme: Hydrolase | 4 |
